# Harnessing artificial intelligence to automate environmental predictions

**DOI:** 10.1101/2025.11.06.684583

**Authors:** Avni Malhotra, Brieanne Forbes, Stefan F. Gary, Amy E. Goldman, Bre Rivera Waterman, Vanessa Garayburu-Caruso, Etienne Fluet-Chouinard, Sushant Mehan, Michael Bruen, Micah Taylor, Marcelo Ardon, M. Bayani Cardenas, Walter K. Dodds, Christian Lønborg, William H. McDowell, Moussa Moustapha, Allison N. Myers-Pigg, Peter Regier, Tod Rubin, Hyun-Seob Song, Ryan D. Stewart, Jorge Villa, Nicholas D. Ward, Timothy D. Scheibe, James C. Stegen

## Abstract

Predicting heterogeneous and non-linear processes remains a fundamental challenge in Earth sciences. Here, we present an artificial intelligence (AI)-guided framework that iteratively combines predictive modeling with targeted field sampling to rapidly improve environmental predictions. We demonstrate our workflow by predicting oxygen consumption, a key process of stream metabolism, across the contiguous United States (CONUS). Our approach consisted of 18 iterative loops of measurements and models, combining distributed participatory field sampling, lab analysis, automated machine learning (ML) predictions, and error and distinctiveness analyses to autonomously guide the next sampling at optimal site locations. Through our approach, we increased the predictive power of sediment oxygen consumption across CONUS by over fifteenfold between the first and last iteration. Relative to our last sampling iteration, our first sampling missed sites with high rates and underestimated median oxygen consumption rates by 68%. In addition to identifying areas of high oxygen consumption rates, iterations enabled refinement of laboratory and data handling methods, and engagement with a broad community of field researchers. We conclude that AI-guided iterative loops between targeted sampling and predictive modeling are a powerful and efficient approach for improving predictions of heterogeneous environmental processes.

## Main text

Predictive modeling of the environment is necessary to understand responses to natural and human-induced stressors, and to support policy and management decisions. However, nonlinear ecosystem processes along with spatiotemporal heterogeneity are major obstacles to a comprehensive predictive understanding of environmental systems^1,2^. Recent developments in artificial intelligence (AI) and machine learning (ML) add a promising new dimension to predictive modeling by expediting the discovery of complex nonlinear processes and modeling heterogeneity across spatiotemporal scales^3–5^. ML models can encapsulate nonlinear interactions and predict process responses at unsampled sites across a wide range of environmental conditions^6–9^. In addition, uncertainty estimations from ML models can be used to determine where future sampling efforts should be focused to efficiently decrease model uncertainty^10,11^. The resulting guidance could help resolve knowledge gaps that impede Earth system process understanding and prediction. Automating the iteration between ML and observations can further expedite discovery, though this AI approach is underexplored in environmental sciences.

Predicting the biogeochemistry of inland waters is one such area that could benefit from AI-guided sampling and modeling. Inland waters are an understudied component of the Earth system that influence human well-being, regulate global biogeochemical cycles, and encapsulate various nonlinear and heterogeneous processes^12–15^. In particular, rivers are major biogeochemical reactors and control points where organic carbon is transformed^16^. Within these transformations, oxygen (O_2_) consumption in riverbed sediments is a fundamental process predominantly fueled by microbial metabolism of organic carbon. This consumption is a key determinant of aquatic energy fluxes and the oxidation-reduction state of sediments, thus controlling the rates and types of biogeochemical fluxes found in river sediments^17^.

The magnitude and variability of O_2_ consumption remain highly uncertain across broad spatial scales, with riverbed sediments estimated to contribute anywhere from 3 to 98% of the total ecosystem (water column plus sediment) O_2_ consumption^18–22^. At local scales, sediment O_2_ consumption is a function of temperature^23^, organic matter and nutrient availability^24^, river size^25^, biota^24^, and hydrology^26^. However, continental to global predictors of O_2_ consumption are difficult to scale from local measurements due to complex interactions among hydrological, geomorphological, and ecological processes^27–30^. The high uncertainty and local-scale drivers of O_2_ consumption in riverbed sediments highlights the need for reduced uncertainty in predictive modeling of such processes.

Here, we present a semi-autonomous iterative approach that combines AI-guided distributed sampling with automated ML modelling to better capture environmental heterogeneity and efficiently improve predictions of biogeochemical processes across continental scales. We demonstrate our approach using the example of river and stream sediment O_2_ consumption rates. Our approach increased the predictive power of O_2_ consumption rates across the contiguous United States (CONUS) over fifteenfold. This workflow greatly improves the efficiency of sampling efforts to capture heterogeneity, leverages the knowledge and resources of local researchers, and showcases the power of AI-driven modeling and distributed, participatory data collection to accelerate scientific discovery and enhancement of predictive capacity.

### A semi-autonomous scientific workflow

The steps to our semi-autonomous approach can be divided into human- and autonomously-driven components (Figure 1 and S1; details in Methods). The human-driven components included distributed field sampling in AI-identified locations, sample processing in the laboratory, data generation, and any necessary revisions of protocols (Figure 1a-c). The autonomous components included ML model training, evaluation, and predictions, and automated error and environmental distinctiveness evaluation to collectively guide the selection of new sites for the next round of field sampling (Figure 1d-f). Thus, sampling was made efficient by targeting sites with high error in the previous sampling-modeling iteration and sites that were environmentally distinct from existing sites. Each stage of the research process was standardized and open access to allow for reproducibility and participation from a global community of researchers.

**Figure 1.**
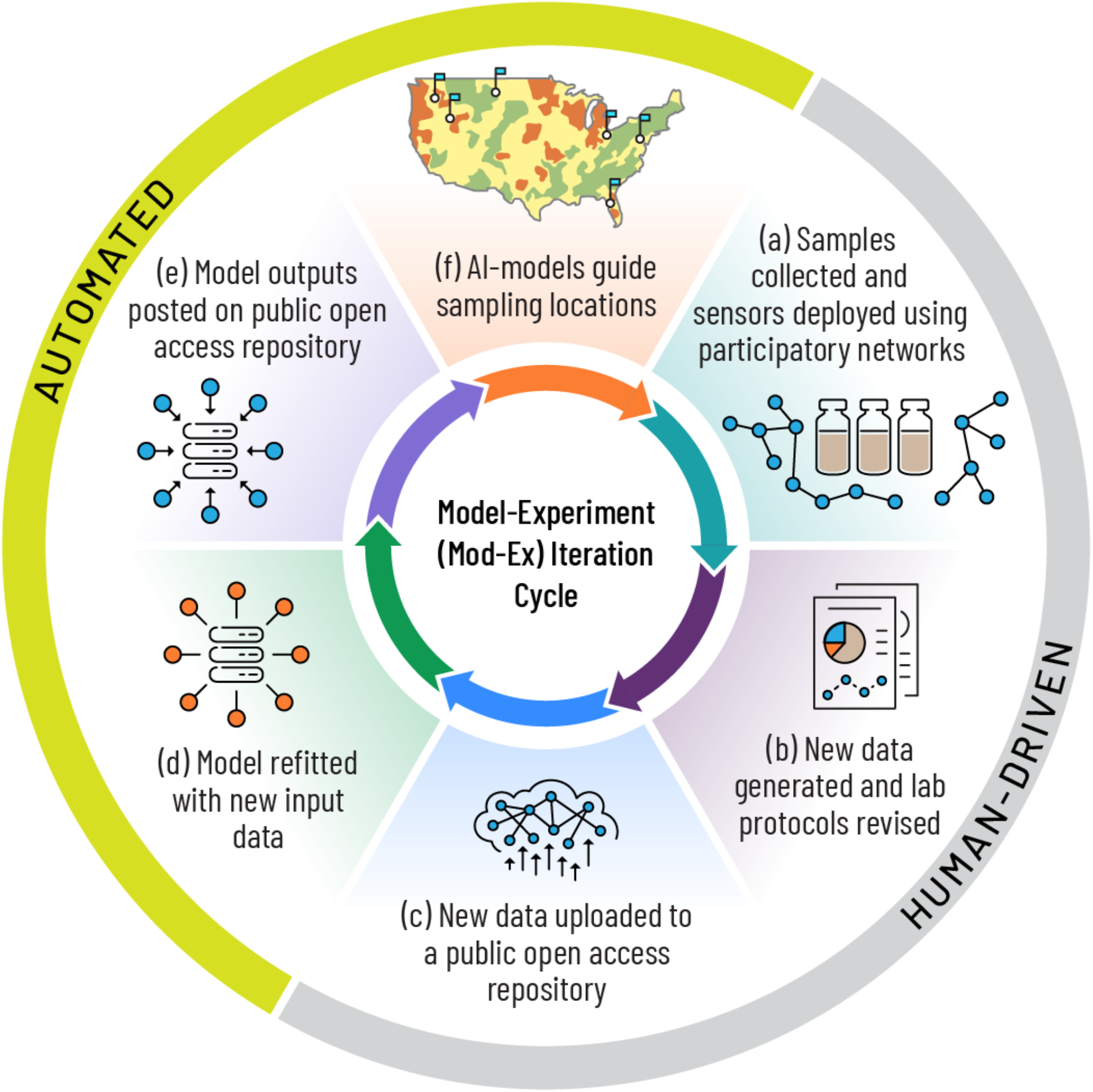
Key components of our semi-autonomous approach that iterates between data generation and modeling multiple times and significantly improves the power to predict an environmental process. We demonstrate our approach using streambed sediment oxygen consumption rates across the contiguous United States (CONUS). (a) sediment samples and sensor data were collected across CONUS via distributed participatory science volunteers, (b) new data were generated from samples using laboratory analyses and any necessary revisions of protocols, (c) generated data were uploaded to an open access data and code repository which triggers the automated generation of (d) new machine learning models, (e) new model outputs were posted on an open-access repository and (f) new model uncertainty and sampling distinctiveness evaluations were used to identify subsequent sampling locations looping back into (a). Iterations can continue until research goals (e.g., level of predictive power) are achieved. A more detailed schematic with specifics of our approach is provided in Figure S1.

The human-driven steps of the workflow were focused on field sampling and laboratory sample processing. We sent standardized sampling kits to collaborators across the CONUS and provided instructions for stream sediment and water sampling. The O_2_ consumption rates were measured in the laboratory using these samples (see Methods). Data generated from field samples were posted to an open-access data repository, which automatically launched the autonomous steps of the workflow. Following data posting, an ensemble of ten stacking-regressor ML models (see Methods) was auto-generated using the new expanded data. For the ML models, we used 25 predictor variables that were derived from in situ measurements and globally-gridded data products (Table S1)^10,31^. The ten ML models at each iteration also helped quantify the uncertainty in model performance. Two types of R^2^ were used to evaluate model performance: 1) predictive power, which was based on unseen test data following training and 2) explanatory power, based on the full dataset (includes training and test data).

At every iteration, ML model prediction error estimates and a distinctiveness evaluation of sampling locations were also generated automatically and informed where the next set of samples should be collected out of a pool of candidate sites (Figure S2). The pool of candidate sites was based on the GLObal RIver CHemistry Database (GLORICH)^32^. The combination of estimated model error at a candidate site and a site’s environmental distinctiveness determined priority levels for the next round of sampling; the more uncertain and distinctive a site, the higher the priority. We balanced our site selection by offering both high and low priority options to our network of distributed participatory collaborators to pick specific sites for sampling. The inclusion of low priority sites helped avoid biasing the models towards the set of environmental conditions that we initially identified as important. Thus, each sampling round was based on a combination of automatically generated site priority and human-coordinated logistics.

The 18 iterations of the project spanned four years, with the first sampling conducted in September 2019 and the last in November 2023. Throughout the iterations, the data and code remained publicly accessible^33^. Furthermore, each step of the research process, from ideation to data generation to analysis and manuscript writing, was conducted in an integrated, coordinated, open, and networked manner (ICON^34^; see Methods) so that collaborators globally could participate in any stage of the research process.

Our approach offered several advantages, including optimizing predictive performance, enabling scrutiny and refinement of methods, and promoting meaningful engagement with site collaborators in a CONUS-scale project. Informing data generation strategies with AI led to more efficient resource use and avoided dedicating effort toward locations that were unlikely to yield significant improvements to predictive performance. The automated prompting of the workflow, combined with the selection of new sampling locations, enabled rapid and flexible iteration. This iterative sampling-modeling framework not only expanded representation across environmentally distinct and undersampled regions but also enhanced the adaptability of our data analysis pipeline and laboratory protocols in response to evolving data characteristics. Thus, while iterative frameworks like adaptive sampling and Bayesian experimental design have been applied to lab and small scale biological research in the past^35,36^, we demonstrate their effectiveness for a continental-scale geoscience application.

### Fifteenfold improvement in predictions of stream oxygen consumption

We found a fifteenfold increase in predictive power of CONUS-scale O_2_ consumption in river sediments between our first and last iterations. Over the course of 18 sampling-modeling iterations, the predictive power (R^2^ based on unseen test data) of ML models increased from less than 0.05 to 0.8 (Figure 2a and Figure S3). Sample size increased from 177 in the first iteration to 684 in the last iteration. Critically, our model improvements with increasing sample size were significantly higher with the highest priority AI-ranked sampling locations when compared to low priority sites (see Figure S3 for a comparison of ML models using low or high priority site data). Thus, simply increasing sample size without targeted sampling would not have increased our predictive power as effectively.

**Figure 2.**
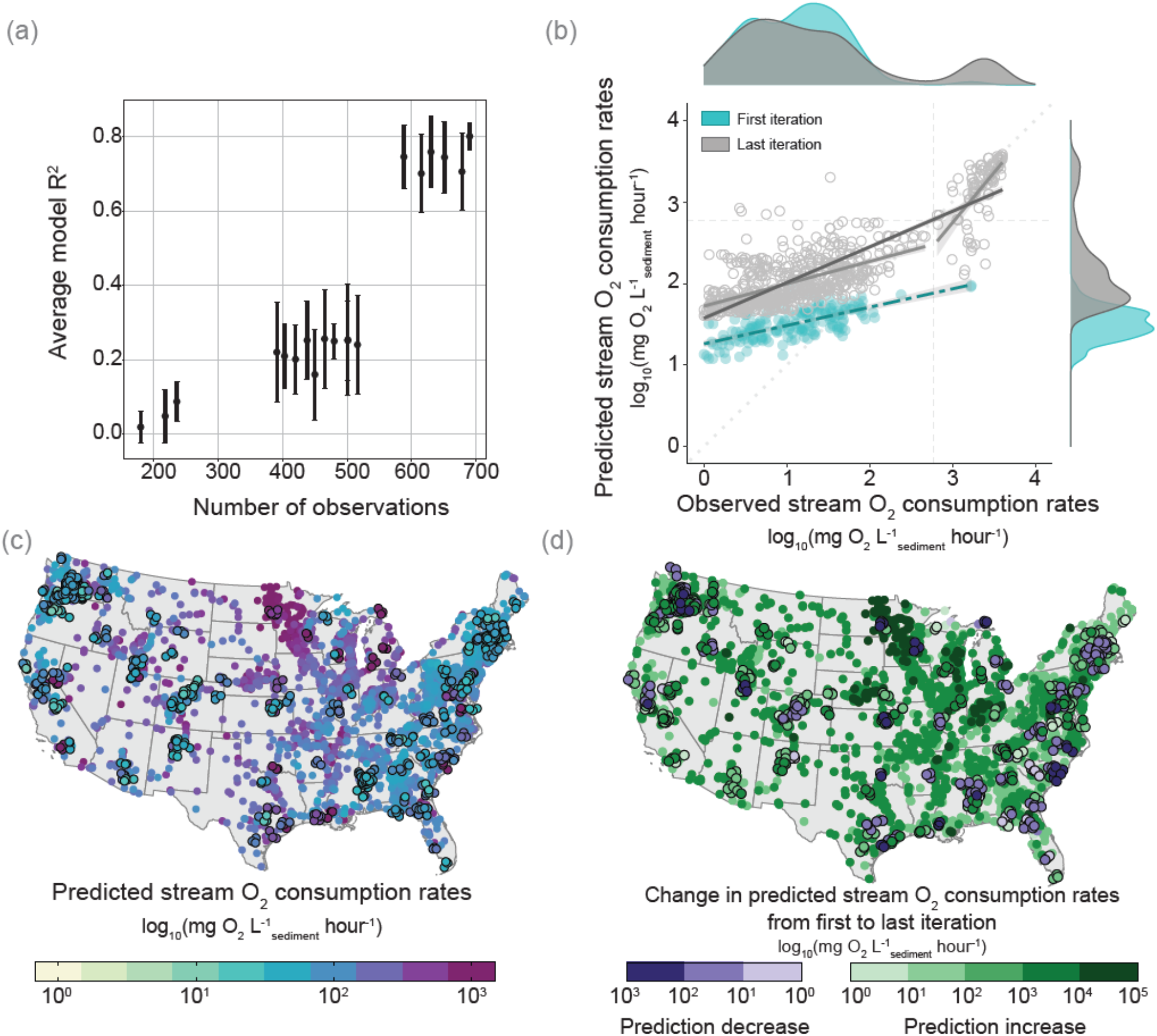
(a) The evolution of the machine learning model R^2^ (predictive power based on test data) with each individual, independent model-sampling iteration. The points and lines show the mean and standard deviation of R^2^ across the ensemble of 10 independent ML stacking-regressor models that were trained for each iteration. (b) Scatterplot with marginal kernel density charts comparing log10-transformed predicted and observed oxygen (O_2_) consumption rates (mean across 10 ML models) from the data at all of the sampled sites. In contrast to panel (a), here R^2^ indicates explanatory power since it is computed with training and testing data combined, which stayed high with R^2^= 0.48 and R^2^= 0.72 for the full dataset after the first (cyan) and the last (grey) iteration, respectively. Raw, untransformed rates of panel (b) are shown in Figure S4. The diagonal light gray dotted line is a 1:1 line. A breakpoint analysis (see Methods) is also shown for the last iteration to better illustrate bimodality of the data. The breakpoint is shown by the vertical and horizontal dashed lines. (c) Predicted O_2_ consumption rates at sampled locations (dots with black outline) and unsampled sites (dots without outline) from the model trained on the last iteration. Unsampled locations are from the Global River Chemistry Database (GLORICH) database^32^ and were used as candidate sampling locations. (d) Change in predicted O_2_ consumption rate between first and last iteration at sampled (dots with black outline) and unsampled (dots without outline) sites.

Furthermore, our iterations allowed us to adjust laboratory and data transformation methods. For example, the large increase in R^2^ between the 12th to the 13th iterations (Figure 2a) is due to revisions in the laboratory methods to account for an increasing number of high O_2_ consumption rates as we gained more samples (Methods). Similarly, we applied log transformation on the data as it became more bimodal with iterations. While predictive power increased (Figure 2a), the explanatory power of our models remained high across iterations (Figure 2b), highlighting that capturing system-wide variability does not necessarily equate to good predictive power at unsampled sites. Across iterations the distribution of O_2_ consumption rates also shifted to a bimodal distribution (Figure 2b) because early iterations largely missed the highest rates. Model performance at the final iteration was particularly strong for both explaining and predicting where high rates should be observed (Figure 2a and 2b), and enabled evaluations of CONUS-scale trends in sediment O_2_ consumption (Figure 2c). Predicted O_2_ consumption rates generally increased first to last iterations (Figure 2d). Notably, relative to our last sampling iteration, rates from our first sampling would have underestimated median CONUS-scale O_2_ consumption rates by 68%. Thus, our approach enabled both increased predictive power (Figure 2a) and better representation of system-wide variability, especially high rates (Figure 2b) across the CONUS.

Our analysis also revealed dominant predictors of O_2_ consumption rates. The final iteration model suggested that variables related to geomorphology, climate, hydrology, land use, and water quality were the strongest predictors of sediment O_2_ consumption rates (Figure 3). Specifically, catchment elevation and slope, water temperature, forest cover, cropland extent, stream water O_2_ concentration and saturation, groundwater table depth, and mean annual air temperature, were the predictors in the final model (dark blue points in Figure 3). In addition to expected macroscale predictors related to climate and hydrology, our results suggest that sediment O_2_ consumption rates are influenced by whether a site is in a mountainous region with little human development or in lower gradient non-mountainous areas with more human development^24,37^. Although mountainous streams can contribute substantially to global carbon cycling, carbon processing in these low dissolved organic carbon systems is more related to rock weathering or turbulence than to biological O_2_ consumption^38^. Conversely, low-lying streams tend to have higher O_2_ consumption rates due to proximity to sources of labile organic carbon coupled with longer residence/contact times with reactive surfaces^39^. Human development, often co-located with streams at lower elevations, can laterally redistribute soil-derived organic carbon from the landscape into aquatic ecosystems^40,41^ and directly impacts sediment O_2_ consumption and overall aquatic ecosystem metabolism^42,43^. Equally important are features associated with agricultural land use. Like other forms of human modification, agricultural practices, such as cover cropping, soil health management, crop rotations, and tillage substantially influence the quantity and composition of organic carbon inputs from the terrestrial landscape to aquatic ecosystems, often enhancing sediment O_2_ consumption rates^40,42,44,45^.

**Figure 3.**
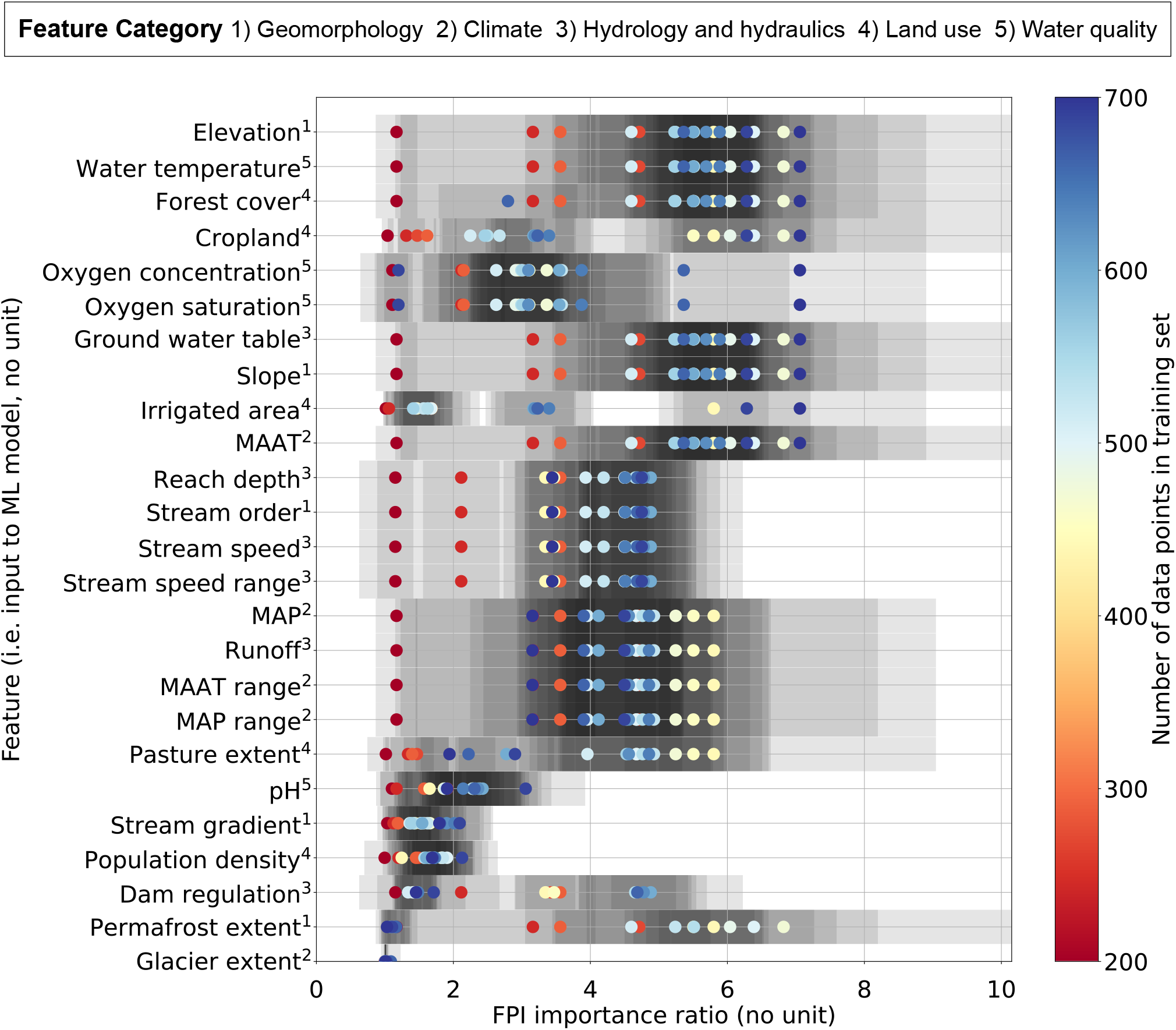
Feature permutation importance (FPI) scores and stability over successive modelling iterations. Machine learning (ML) model inputs (i.e. the list of features on the vertical axis) are grouped into five categories: geomorphology, climate, hydrology and hydraulics, land use, and water quality. The mean FPI scores for each iteration are plotted as dots whose color corresponds to the number of data points in the training data for that iteration. The uncertainty of each FPI data point, as quantified by the standard deviation of the FPI scores over the ML model ensemble members for each iteration, is represented by a translucent gray box. Each FPI data point has a unique uncertainty range box with the same level of translucency; the darker grays correspond to regions where one or more of the FPI data points’ uncertainty range boxes overlap. Darker grays signify more overlapping uncertainty ranges than lighter grays. Variables ordered together on the y-axis are correlated to each other.

Iterations in our approach also allowed us to understand how predictor importances, i.e., feature permutations importance (FPI), changed and persisted over iterations (Figure 3 and Figure S5). Future users of our workflow could decide to continue iterating until they reach a desired R^2^ or until they reach stability in predictor FPI. In our case, the R^2^ plateaued in the last six iterations (Figure 2a), but FPI continued to vary across iterations (light to dark blue points in Figure 3). Dominant predictors of sediment O_2_ consumption varied across the 18 iterations and predictors varied in whether their scores increased or decreased over time (Figure S5). For example, while the predictor importance of pH never increased, the predictor importance of mean annual precipitation increased in early iterations but decreased in the final iterations (Figure S5). As the dataset increased with an ever-widening range of values for each predictor variable, it is likely that the correlations between variables changed, and thus, the composition of dominant predictor groups changed. Thus, the FPI for each group of variables may change for successive iterations as new samples provide more information about the influence (or lack of influence) of each variable. Our approach emphasizes that the traditional approach of one sampling and modeling iteration could provide predictors that are very different from predictors after multiple iterations, thereby skewing conceptual understanding of system function.

### Benefits and challenges of the distributed and participatory sampling approach

The participation of distributed volunteer collaborators was critical to the human-driven components of our workflow but came with a variety of challenges such as standardization, logistical challenges, and engaging participants. We maximized consistency using standardized kits, written and video protocols, and by analyzing samples in one analytical facility. While consistency in sampling and analysis was a clear strength of our approach, it required additional effort to ensure consistent protocols and interoperability across iterations. There were also complex logistics related to shipping and receiving sampling kits from across the CONUS, an activity we conducted multiple times a month for the duration of the project. Another challenge was finding sufficient numbers of people through time, keeping them engaged, and finding people in new priority locations. We overcame this challenge by repeatedly announcing opportunities to sample and through targeted searches for new participants. Thus, we acknowledge the “semi” in our semi-autonomous approach wherein the sampling components required considerable human intervention, even if the modeling was fully automated.

Despite challenges, human involvement from a broad community of collaborators greatly benefited the study before, during, and after sampling. The human-driven components of the workflow were also critical for method refinement. For example, we learned early on that data were bimodal and were able to adapt lab methods to better measure high O_2_ consumption rates. Over the iterations, we engaged with hundreds of participants, which included approximately 75 individuals who regularly joined meetings, 130 who conducted sampling, 10 who conducted analyses, and 25 who helped with manuscript writing. Broad engagement via participatory science enabled efficiency gains in distributing sampling across volunteers and improved study design from a wide range of perspectives. We emphasized accessibility through open data and code, and co-developed standards for psychological and physical field safety. The project also led to new networks and connections among our participants, including numerous early-career researchers.

## Conclusions

We demonstrate a scientific workflow of rapid iterations between AI-guided sampling and autonomous modeling of environmental systems that can be applied to any dynamic and heterogeneous system. As a case study, using this approach, we observed a fifteenfold improvement in model predictions of CONUS-scale sediment O_2_ consumption rates. Our approach provides rich opportunities for a deeper exploration of drivers influencing model performance and supports the targeted refinement of data-driven hypotheses. We conclude that leveraging AI and automation capabilities, coupled with traditional ML and human-driven distributed sampling, can accelerate scientific discovery and improve the fidelity of predictive models. While applicable to a variety of domains, our approach is effective for environmental systems where process complexity, non-linearity and spatiotemporal heterogeneity present persistent challenges to predictive modeling. Particularly promising are contexts where broad-scale predictor data are available, sample analysis and collection have a feasible turnaround time, and an accessible field network already exists. Our approach is especially valuable in contexts where fine-scale drivers must be resolved to inform regional or global-scale predictions critical for resource management and policy, for example, groundwater pollution, disturbances like drought or wildfire, biodiversity, forest biomass, and human health. In addition to predictive power, our approach provides numerous benefits for efficient resource allocation, open access, reproducible science, and collaborative opportunities that are especially vital for early career researchers.

## Methods

### Overview

Below, we detail each step of our measurement and modeling loops (Figure 1). The main steps include field sampling planning, sampling of river and stream sediments, laboratory quantification of O_2_ consumption, machine learning (ML) model implementation, and error and distinctiveness quantification to guide subsequent field sampling^46^. These sampling-modeling iterative loops were conducted 18 times between September 2019 and November 2023.

Throughout the paper, we use artificial intelligence (AI) as an umbrella term that includes ML models but also the automated use of ML model outcomes to generate guidance on ideal sampling locations. Conversely, we use the term ML when specifically referring to the ML models used in our approach. Thus, the AI in our approach consists of automation using human-defined rules on how ML outcomes should inform future sampling. While the ML in our approach is traditional ML wherein rules are being learned from the data.

### Prior to field sampling

Community discussions were held prior to the first sampling in September 2019. Additionally, two virtual open discussions were held in December 2021 and February 2022 to present further study design, discuss questions, and receive feedback from the broader scientific community. The open discussions were widely advertised on message boards, mailing lists, social media, conference presentations, and through direct communications. The open discussion calls had over 50 attendees each, and the recordings were made public on YouTube for people to watch and provide feedback asynchronously. The study design was modified based on attendee and asynchronous feedback with the aim of improving scientific impact and enhancing mutual benefit with the broader communities engaged.

### Field sampling

The opportunity for volunteers to engage in sampling and data analysis was widely advertised via direct emails, listserv announcements, and conferences. From April 2022 to October 2023, volunteers were sent sampling kits to collect sediment and stream water. Sampling locations were selected following predictions from the ML models described in more detail below. In addition to sample collection, volunteers also deployed a dissolved oxygen and temperature sensor, and measured pH. A detailed description of sampling protocols, field metadata, and site contextual photos were published along with the source data ^33,47^. A video protocol was also made available on YouTube (https://tinyurl.com/CM-video-protocol). Briefly, shallow subsurface sediments (1-3 cm) were collected from depositional zones and sieved in the field to <2 mm into 500 ml brown Nalgene bottles. Additionally, 500 ml of unfiltered surface water was collected at each site. Both sediments and water were kept on blue ice while they were shipped to the Pacific Northwest National Laboratory (PNNL) campus in Richland, Washington (USA) within 24 hours of collection.

Between the third and fourth iteration (Figure 2a), there is a rapid increase in the number of sampled locations due to an additional sampling campaign in the Yakima River basin (YRB) in Pacific Northwest USA. Here, the methods to guide sampling were different but the sampling methodology was the same as the other samples in this study. Briefly, the criteria for sample location selection for YRB were based on variable importances for a process model ^48^ where a random forests analysis was used to map the input and output of the model. This resulted in spatial clusters across the YRB which were used to guide the sampled locations ^49^. The sampled locations were distributed across the clusters and representative of the larger Columbia River basin region. The sampling and laboratory methods were consistent with the rest of the samples ^50^.

In total, we had 185 unique sites from across the CONUS and 47 from YRB. For each unique site, we had 3 replicates, resulting in a total of 684 observations (12 samples were lost).

### Laboratory analyses

Once in the laboratory, the sediments were sieved again to <2 mm and homogenized. About 10 mL (to account for slow O_2_ consumption) and another 2.5 mL (to accommodate fast O_2_ consumption) of sediments were then subsampled into separate 40 mL glass vials (I-Chem amber VOA glass vials) containing a 0.5 cm diameter pre-calibrated oxygen sensor (PreSens GmbH, Regensburg, Germany). After subsampling was completed, vials with sediment were covered with a Breathe-EASY membrane to allow airflow. Further, unfiltered stream water and sediment vials with Breathe-EASY membranes were placed on their side to maximize exposure to O_2_ and left in the dark at 21°C the night before starting the experiments to acclimate to a standard temperature.

O_2_ consumption measurements were performed following previously employed methods^51,52^. Briefly, unfiltered stream water was added to the sediments, filling up the vial halfway and agitating vigorously to aerate the sediment and water mixture. The vial was then filled completely with unfiltered stream water to minimize headspace and a time zero dissolved O_2_ (DO) measurement was recorded. The vials were then placed horizontally on an orbital shaker at 250 rpm for 2 hours. DO was measured noninvasively every 5 minutes for the first 30 minutes and every 30 minutes after that using an O_2_ optical meter (Fibox 3; PreSens GmbH, Germany). O_2_ consumption rates were calculated as the slope of the linear regression between DO concentration and incubation time for each reactor. Whenever O_2_ consumption was slow, the measurements from the 10 mL incubations were used to calculate rates. If O_2_ consumption was fast (i.e., DO was <= 5 mg L^-1^ at time zero in the 10 mL incubations), measurements from the 2.5 mL incubations were used as they captured O_2_ dynamics in each reactor with higher resolution than the 10 mL incubations, which went anaerobic too quickly to reliably estimate an O_2_ consumption rate. Initially, all O_2_ consumption rates were calculated using 10 mL sediment incubations and thus they were not normalized because the ratios of sediment and water were the same across samples (i.e., mg O_2_ L^-1^ h^-1^). As the breadth of sampled environments increased across model-experiment iterations, we encountered faster O_2_ consumption rates (Figure 2b), leading to the introduction of 2.5 mL incubations. Once this occurred, we began normalizing the rates by the volume of sediment in the vial to account for variation in the ratios between sediment and water across samples and enable cross-sample comparability^53^.

### Machine learning model input features

Measured O_2_ consumption rates were then modeled using ML model ensembles. The ML models discussed in this manuscript all have 25 inputs, or features, which were used to predict O_2_ consumption rates (see Table S1 for a list of features). Four of these features – temperature, DO in mg L^-1^, DO in percent saturation, and pH – were *in situ* observations taken during sample collection. The remaining 21 features were broad-scale indices derived from the RiverAltas global database^54^. We subdivided RiverAtlas into tiles for faster access during subsequent processing. We then found the closest river segment in RiverAtlas for each field site and used the properties of that RiverAtlas segment as the additional features describing each field site. This data fusion process occurred at the 30-m spatial resolution of RiverAtlas. RiverAtlas contains more than 100 features; we selected 21 features based on qualitative practical considerations, scientific relevance, and the desire to reduce the number of correlated features so as to span a wide range of environmental contexts but not have excessive amounts of duplicated information. To make predictions at sites where *in situ* observations were unavailable, we relied on the Global River Chemistry Database (GLORICH) database^32^. Lastly, we ruled out the influence of the timing of sampling on our model predictions by confirming that hour and month had no relationship with O_2_ consumption predictions (Figure S6 and S7).

### Machine learning models

All ML models trained in this work were run using the scikit-learn framework^55^, using the ensemble of ensembles approach of Gary et al^10^. Each iteration of the ML workflow consisted of an ensemble of ten ML stacking-regressor models independently trained on different, randomly selected training/testing data splits of the same initial data. All of the randomized data splits are 75% for the training set and 25% for the testing set. This first layer of ensembles was an implementation of Monte-Carlo cross-validation^56^ for a given training dataset; the spread in the ensemble members’ scores was the basis for the uncertainty ranges (Figures 2a and 3). For a given iteration, final O_2_ consumption rate predictions were the mean of the individual predictions over all ten of the cross-validation ensemble members at each site. Each ML model in the cross-validation ensemble was a stacking-regressor because it is, in turn, an ensemble of 15 “submodels” that were trained in parallel and whose predictions were combined as the weighted average of the submodels. Submodel weights were determined by finding the weights that minimize the combined prediction error on the training data using a non-negative least squares algorithm^31^. Stacking-regressor ML models allow different types of ML algorithms to be blended into a single ML model and typically result in models whose scores are equal to or better than the best performing single submodel in the stacked ensemble^57^. In this case, each of the 15 submodels is a different ML algorithm as described in Gary et al^9^. In addition to independently fitting the submodels in each ML model, the ML workflow also optimized all submodel hyperparameters in parallel^10^.

### Breakpoint analysis

A nonlinear relationship of predicted versus observed O_2_ consumption rates emerged as we progressed across model-experiment iterations (Figure 2b). This prompted us to conduct a breakpoint analysis to identify how to split the data into low and high O_2_ consumption rate groups. To do so, we ran a piecewise linear regression to identify the statistical breakpoint using the ‘segmented’ package in R^58^. The analysis was performed for each model iteration, with the final breakpoint ranges shown in Figure 2b and the change in the breakpoint over time/model iterations shown in SI GIF1. The data points above the breakpoint were classified as ‘hot spots’, or samples with high rates of O_2_ consumption, while points below this threshold were classified as ‘cold spots.’ Despite most of the initial data samples being from cold spots, the strategy for selecting new sampling sites was, as described below, weighted toward candidate sites that were predicted to be hot spots thus ensuring a substantial group of likely hot spots above the breakpoint were selected in subsequent iterations, increasing the range of behaviours sampled (Figure 2b).

### Guiding sampling locations using model uncertainty and environmental distinctiveness

There were two key innovations of this work that facilitated open and semi-autonomous research. First, the modeling component of each model-experiment iteration was fully automated as soon as data were published with near real time results available publicly. Second, each iteration provided high priority sites for the next iteration. The first of these innovations was implemented using GitHub Actions to launch our ML workflow when the training data, posted publicly on GitHub, were updated with new data. In our case, the GitHub Action uses an application programming interface (API) call to the Parallel Works ACTIVATE compute platform^10^ to launch the workflow on a cloud cluster. The process by which the ML workflow predicts the highest and lowest priority sites was based on the combination of two sets of information generated for each GLORICH candidate site during the ML model evaluation process: 1) an estimate of the ML model prediction error at each GLORICH site and 2) a metric of how distinctive the environmental conditions (based on the 25 ML input features) of a particular GLORICH site were, relative to already sampled sites.

First, we used a linear regression to predict the error of ML predicted O_2_ consumption rates at a candidate GLORICH site. This is because while it is possible to quantify the ML models prediction error at the sampled sites by finding the difference between the observed and predicted O_2_ consumption rates, this was impossible at GLORICH candidate sites where we did not have observed O_2_ consumption. Thus, we conducted a linear regression between the ML predicted O_2_ consumption rate and the observed error at all of our sampled sites. Using this regression function and predicted O_2_ consumption rates at GLORICH sites, we predicted a model error for each GLORICH site as the 95% confidence interval for the mean value of the ML prediction error. As such, the predicted errors of the ML model change at each site with each iteration and are proportional to the magnitude of the predicted O_2_ consumption rates. During the course of the iterations, we tested a second method for estimating the error of ML model predictions that used the ML model ensemble spread of the predicted O_2_ consumption rate at a particular site; while this method was better at estimating the error, the magnitude of the error was still proportional to the predicted O_2_ consumption rate (not shown) so we kept using the initial method described above.

Second, our site priority score included a metric for the distinctiveness of a potential site’s environmental context compared to all other observed and candidate sites. To define the distinctiveness of a site’s environmental context we first projected all previously sampled and potential sites onto the first two principal component axes, derived from a principal component analysis (PCA) using the geospatial variables described above. A potential site’s distinctiveness was estimated as the Euclidean distance from that site to the centroid of the sites that had already been sampled. Larger distances were interpreted as indicating greater distinctiveness (Figure S2). The PCA tools implemented in scikit-learn were used for this analysis^55^.

Finally, the two metrics of ML model error and environmental distinctiveness were combined as a final site priority score. This was done by an unweighted multiplication of the ML estimated error and PCA-derived environmental distinctiveness, each normalized by its respective maximum value. This provided priority scores for the next field sampling iteration. We added the top 10% and bottom 10% of priority scores to a public map and asked volunteers to choose locations on the map to sample. The bottom 10% were included to avoid strongly biasing new data, while the top 10% was intended to challenge the model and expand its transferability across divergent environments. If a collaborator was interested in sampling somewhere that was not on the map, we provided them the opportunity to collect some preliminary data (i.e., pH, temperature, and DO) to input into the model and determine priority in the same way as other sites. Every month, after the sampling, field, and lab methods described above, laboratory-generated O_2_ consumption rates were combined with field (meta)data and pushed to GitHub. Using a GitHub Action, when a release was created, model training automatically started running on the cloud cluster^10^. All models and results were automatically posted to GitHub. The ensemble ML model was rebuilt in this way every month, producing a new set of predictions and prioritized sites. An updated map was also provided for collaborators to volunteer to sample. These iterations resulted in AI-guided continental-scale sampling integrated with model testing and improvement, on a monthly basis, for a total of 18 model-experiment cycles.

### Evaluating the role of AI-guided sampling in increasing model performance

Since our sample size also increased with iterations (which would increase model R^2^), we evaluated the extent to which our AI-guided site selection impacted the progressive improvement of model performance. To do this, we ran the ML workflow as *in silico* experiments using observed O_2_ consumption rates. We ran the workflow six times using only the 100, 200, and 300 highest and lowest priority samples in the dataset as determined by the last iteration training data (HP and LP points in Figure S3). ML model performance was significantly higher when trained on only the highest priority sites compared to lowest priority sites. Thus, the AI-guided site selection likely accelerated the increase in model performance relative to random, unguided sampling. We are unaware of other methods to clearly evaluate how much faster the model improved due to AI-guided site selection. Using a random selection of sites to perform an *in silico* experiment would not be sufficient because that would require use of our observed data, which were not derived from random sampling. Our observations are AI-guided such that we cannot use them to generate a true random-sampling expectation. Doing so would require a parallel study using only randomly selected field sites, which was unfeasible. Future studies could gain additional insights through a digital twin approach in which an *in silico* version of CONUS is set up with variation in O_2_ consumption rates as a function of environmental variables. That digital twin could be sampled through random and AI-guided methods to provide a truly fair comparison. We encourage development of such digital twin capabilities so that future studies can evaluate the value added by using an AI-guided approach prior to implementing a given study.

## Data availability

Data and code are available on the Environmental System Science Data Infrastructure for a Virtual Ecosystem Repository (ESS-DIVE) at https://data.ess-dive.lbl.gov/datasets/doi:10.15485/2998468.

## Acknowledgements

This research was supported by the U.S. Department of Energy (DOE), Office of Biological and Environmental Research (BER), Environmental System Science (ESS) Program as part of the River Corridor Science Focus Area (SFA) at the Pacific Northwest National Laboratory (PNNL). PNNL is operated by Battelle Memorial Institute for the DOE under Contract No. DE-AC05-76RL01830.

