## Supplementary Information for "Harnessing artificial intelligence to automate environmental predictions"

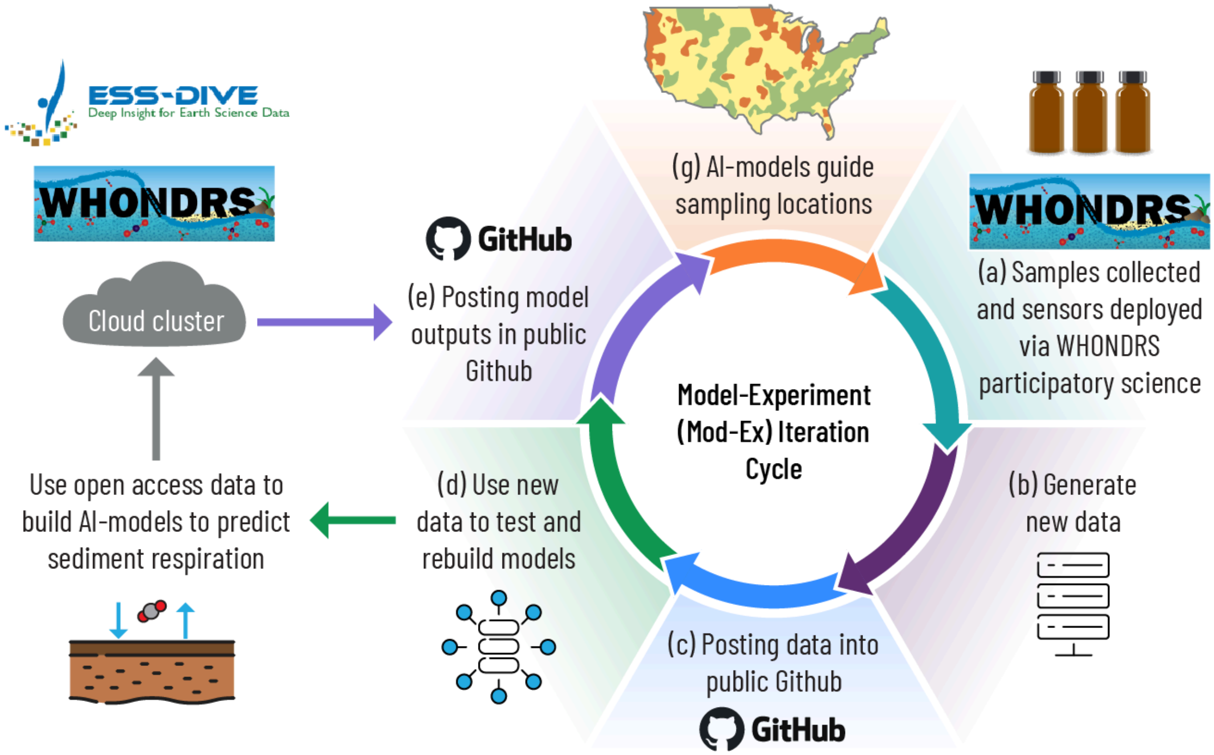


**Figure S1**. Key components of our semi-autonomous approach that iterates between data generation and modeling. (a) sediment samples and sensor data are collected across contiguous United States (CONUS) via distributed participatory science volunteers via the Worldwide Hydrobiogeochemistry Observation Network for Dynamic River Systems (WHONDRS), (b) samples are thereafter analyzed in the laboratory, (c) generated data are uploaded to a data and code repository (in our case GitHub) which triggers the automated generation of (d) new machine learning models, (e) new model outputs are posted on GitHub and (f) new model uncertainty and sampling representativeness analyses are used to identify subsequent sampling locations looping back into (a).

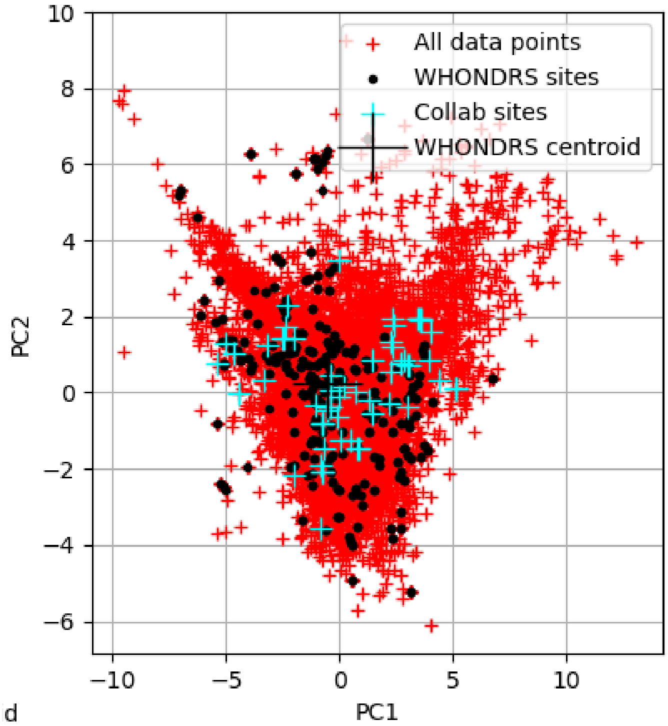


**Figure S2**. Example of the principal component analysis (PCA) results quantifying the distinctiveness of the environmental context of sampled and candidate sites. This figure was autogenerated during the running of the first machine learning (ML) ensemble member of the last iteration. The magnitude of each site's two principal components with the greatest variance ("PC1" and "PC2") are plotted with respect to each other. Red and cyan data points correspond to candidate sites where there are no oxygen consumption observations while black dots correspond to sites where sediment samples had been already collected at the time of this particular iteration. The cyan points are sites where members of our sampling community (WHONDRS) expressed interest in sampling and have the ability to collect sediment samples while the red points are the locations of all possible candidate sites. Each site's environmental distinctiveness is quantified by its distance from the centroid of the whole dataset in this two dimensional plot. It is a subset of the cyan points that were selected for the next iteration of the ML workflow contingent on combining each centroidal distance visualized here with the estimated model error at that point and real-world sampling logistics.

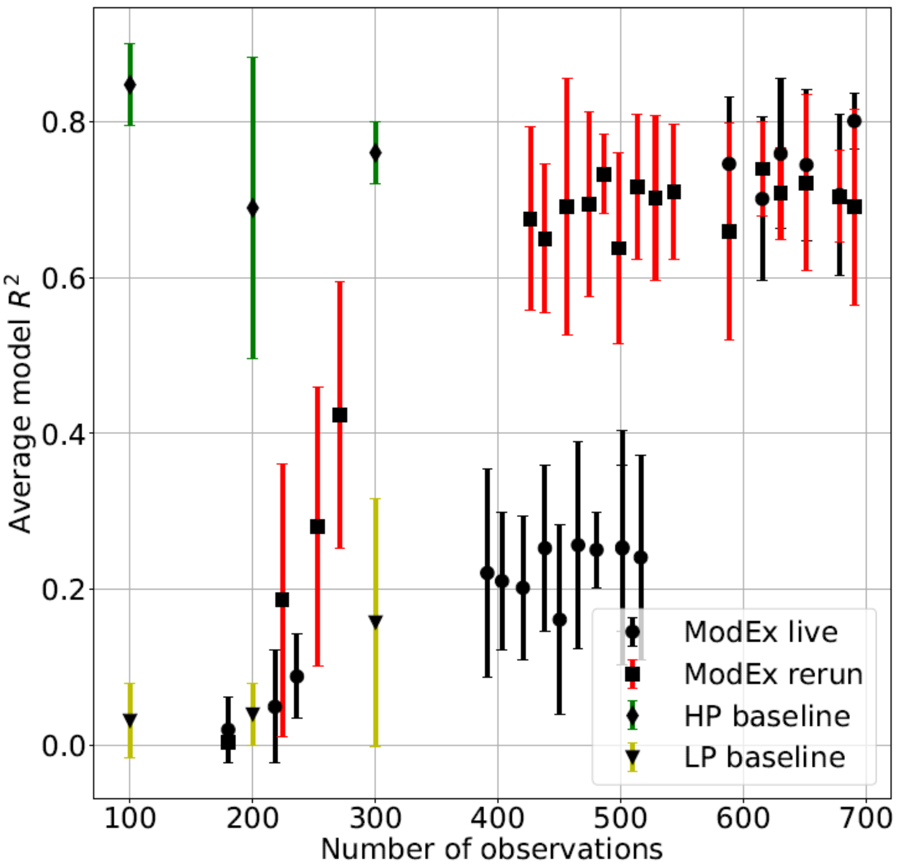


**Figure S3**. The evolution of the machine learning (ML) model $R^2$ as each model-sampling iteration was completed. The "live" iterations are the models that were automatically generated at each loop and that were used for making decisions on which sites to sample in the next loop. The "rerun" iterations were conducted at the end of the project and use a $\log_{10}$ filter because this reduces the bias of the ML models. The red, black, green, or yellow uncertainty envelopes are one standard deviation of the scores of the 10 ML model ensemble that was generated for each iteration. We also conducted a retrospective evaluation of how the iterations might have progressed if all the data had been available at the beginning, and if sample size had been increased using only the lowest priority sites (LP baseline) or only the highest priority sites (HP baseline). HP having much greater predictive power than LP demonstrates that using the AI-guided sampling greatly improved model performance.

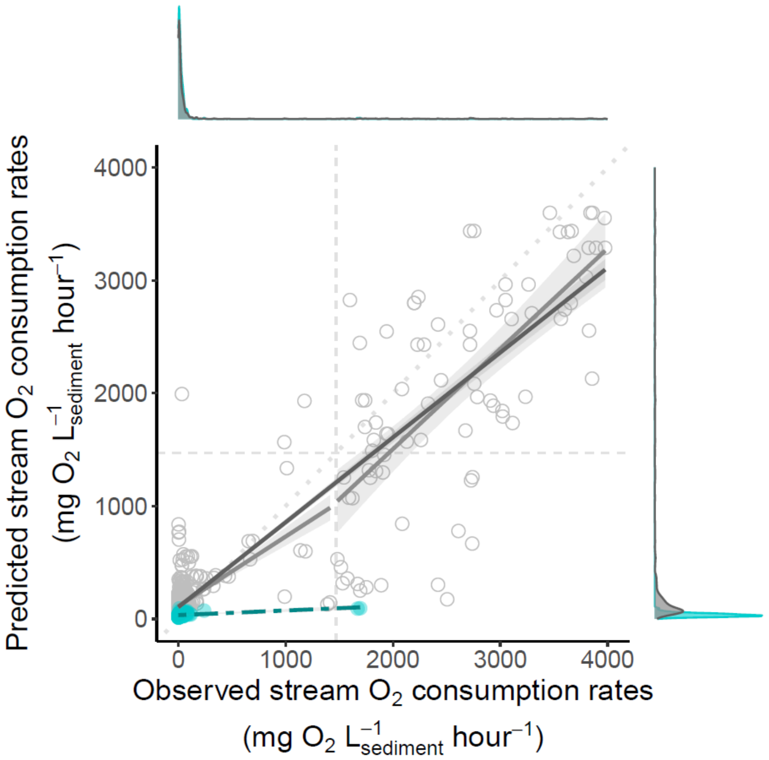


**Figure S4**. Scatterplot with marginal kernel density charts comparing predicted and observed oxygen ($O_2$) consumption rates (mean across 10 machine learning models) from the data at all of the available sites; here, $R^2$ indicates explanatory power, with $R^2$= 0.22 and $R^2$= 0.85 for first (cyan and dashed) and last (dark grey and solid) iteration, respectively. The first iteration is difficult to see due to the distribution of the data and can be better seen in Figure 2b. Dotted line is a 1:1 line. A breakpoint analysis (light grey) (see Methods) is also shown for the last iteration to better illustrate bimodality of the data.

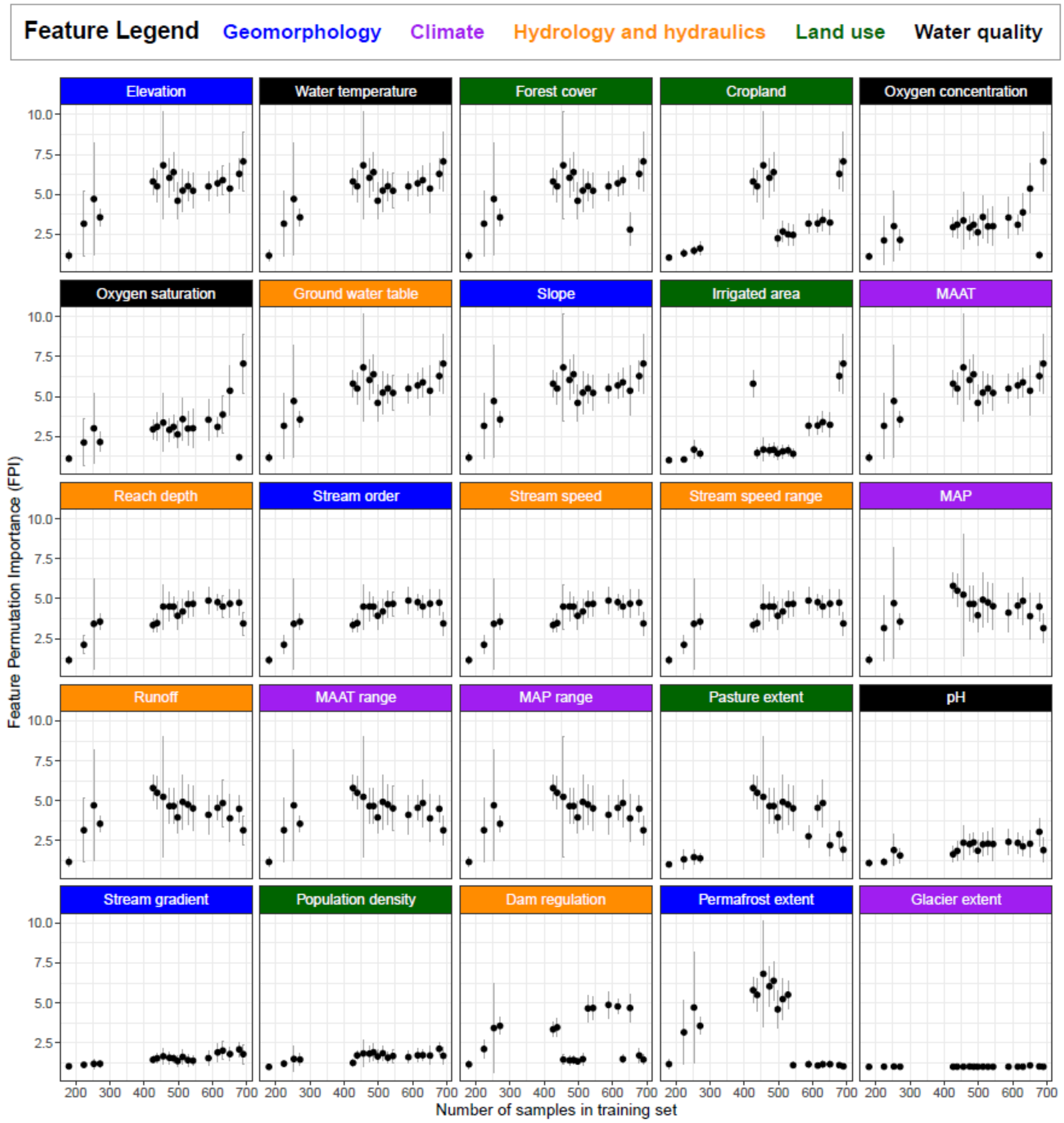


**Figure S5.** Feature permutation importance (FPI) of each feature over each iteration.The grey error bars indicate the standard deviation of the FPI scores over the ten 10 cross-validation machine learning (ML) model ensemble members for each iteration. The order of features is determined by the last iteration (i.e. the most important variables in the last iteration are in the top left and the least important are in the bottom right). The features are grouped and color-coded into the 5 categories of geomorphology (blue), hydrology and hydraulics (orange), water quality (black), climate (purple), and land use (green). Our ML approach allows us to evaluate changes in FPI importance over time. We acknowledge that sharp changes in data patterns can occur due to the inherent dynamics of feature correlations and sample distributions across iterations. These fluctuations are expected and can be indicative of varying feature interdependencies and data sample characteristics. The shading in Figure 3 captures the range of possible importance scores, offering a probabilistic perspective on feature significance rather than relying solely on fixed data points. This method provides detailed insights into feature importance dynamics, enabling comprehensive understanding of predictive model behavior over iterative sample inclusion.

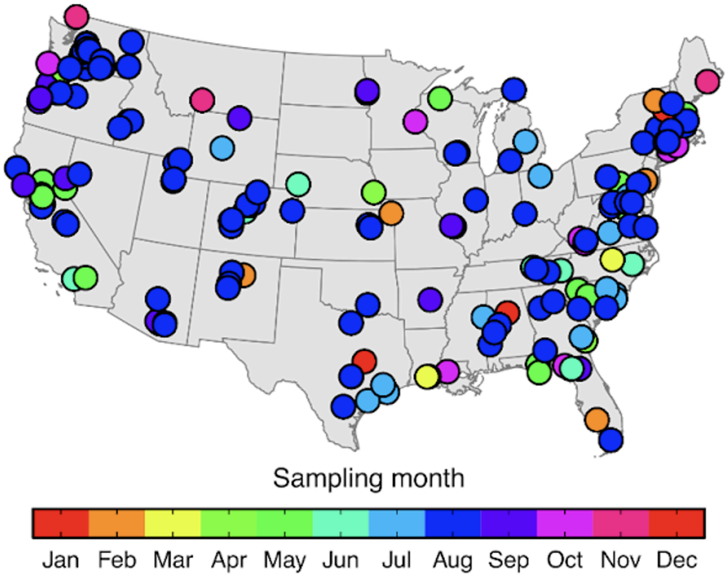


**Figure S6.** Map of sampled sites across the contiguous United States (CONUS) with the color in-fill representing the calendar month when samples were taken.

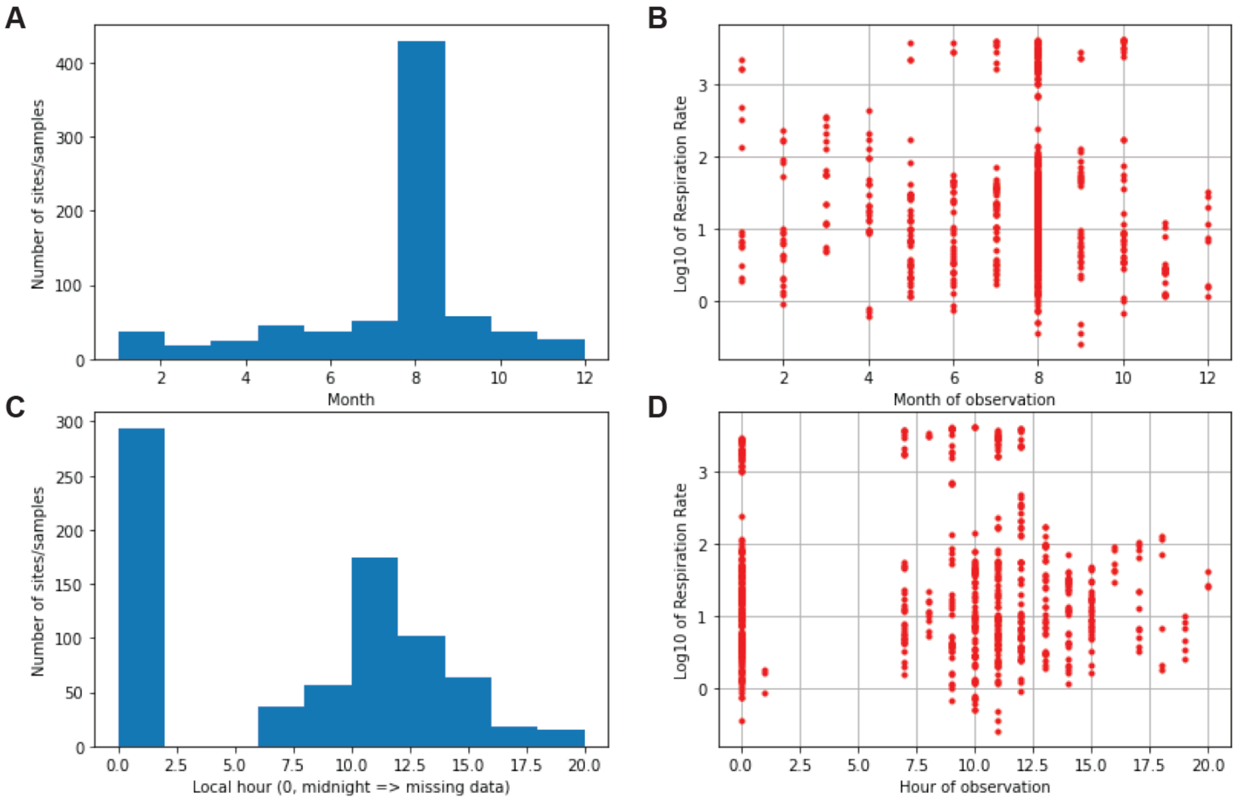


**Figure S7**. (a) Distribution of samples collected by month. (b) Distribution of predicted oxygen ($O_2$) consumption rates by month of sampling showing no overall trend. (c) Distribution of local hour of sampling. (d) Distribution of predicted $O_2$ consumption rates by hour of sampling; which showed a slight trend with predicted $O_2$ consumption rates which we accounted for by using mean in situ temperature as a predictor.

**Table S1**. Summary of the feature permutation importance (FPI) results across the model-sampling iterations. The table presents the feature names and detailed descriptions corresponding to the feature identification numbers used for plotting. Features with exactly the same FPI scores and uncertainty range for any particular iteration were correlated with $R > 0.5$ for that iteration and were thus permuted together as a block to prevent biasing FPI scores with the "leakage" of information (i.e., information from the validation dataset influences the training process) from one correlated feature to another. WHONDRS sourced data were community collected at their sites. RiverAtlas data were downloaded on November 20th, 2020.

| Predictor short name | FPI in last iteration | Standard deviation of FPI | Uncertainty in importance | Source | Unit | Definition |
|---|---|---|---|---|---|---|
| ele_mt_cav | 7.06 | 1.85 | 10.86 | RiverAtlas | m | Mean elevation over reach catchment |
| Mean_Temp_Deg_C | 7.06 | 1.85 | 10.86 | WHONDRS | degrees_Celsius | Mean water temperature from dissolved oxygen (DO) sensor (miniDOT) in degrees Celsius measured during in situ sensor deployment. |
| for_pc_cse | 7.06 | 1.85 | 10.86 | RiverAtlas | percent | Forest cover extent over reach catchment |
| crp_pc_cse | 7.06 | 1.85 | 10.86 | RiverAtlas | percent | Cropland extent over reach catchment |
| Mean_DO_mg_per_L | 7.06 | 1.85 | 10.86 | WHONDRS | milligrams_per_liter | Dissolved oxygen measured during in situ sensor deployment. |
| Mean_DO_percent_saturation | 7.06 | 1.85 | 10.86 | WHONDRS | percent_saturation | Dissolved oxygen saturation measured during in situ sensor deployment. |
| gwt_cm_cav | 7.06 | 1.85 | 10.86 | RiverAtlas | cm | Mean ground water table depth over reach catchment |
| slp_dg_cav | 7.06 | 1.85 | 10.86 | RiverAtlas | degx10 | Mean terrain slope over reach catchment |
| ire_pc_cse | 7.06 | 1.85 | 10.86 | RiverAtlas | percent | Irrigated area extent (Equipped) over reach catchment |
| tmp_dc_cyr | 7.06 | 1.85 | 10.86 | RiverAtlas | deg. C | Annual avg. air temp. over reach catchment |
| RA_dm | 3.44 | 0.71 | 4.67 | RiverAtlas | m | Est. avg. depth of reach derived from other RA vars. |

| | | | | | | |
|---|---|---|---|---|---|---|
| RA_SO | 3.44 | 0.71 | 4.67 | RiverAtlas | non-dim. | Stream order |
| RA_ms_av | 3.44 | 0.71 | 4.67 | RiverAtlas | m/s | Annual avg. stream speed |
| RA_ms_di | 3.44 | 0.71 | 4.67 | RiverAtlas | m/s | Range between annual min and max avg. stream speed |
| pre_mm_cyr | 3.16 | 0.91 | 4.68 | RiverAtlas | mm | Annual avg. precipitation over reach catchment |
| run_mm_cyr | 3.16 | 0.91 | 4.68 | RiverAtlas | mm/year | Annual avg. land surface runoff over reach catchment |
| tmp_dc_cdi | 3.16 | 0.91 | 4.68 | RiverAtlas | deg. C | Range of annual min and max air temp. over reach catchment derived from RA data |
| pre_mm_cdi | 3.16 | 0.91 | 4.68 | RiverAtlas | mm | Range of annual min and max precip. over reach catchment derived from RA data |
| pst_pc_cse | 1.94 | 0.67 | 3.63 | RiverAtlas | percent | Pasture extent over reach catchment |
| pH | 1.91 | 0.76 | 3.87 | WHONDRS | pH | pH |
| sgr_dk_rav | 1.80 | 0.59 | 3.00 | RiverAtlas | dm/km | Avg. stream gradient over reach |
| ppd_pk_cav | 1.69 | 0.50 | 2.42 | RiverAtlas | people/km^2 | Average population density over reach catchment |
| dor_pc_pva | 1.45 | 0.31 | 2.22 | RiverAtlas | percent | Degree of dam regulation on reach |
| prm_pc_cse | 1.03 | 0.06 | 1.15 | RiverAtlas | percent | Permafrost extent over reach catchment |
| gla_pc_cse | 1.01 | 0.01 | 1.03 | RiverAtlas | percent | Glacier extent over reach catchment |

**Supplementary GIF1: break point analysis over iterations**
https://github.com/WHONDRS-Hub/ICON-ModEx_Open_Manuscript/blob/main/fig-model-score-evolution/scatter.gif
